# Concurrent cognitive, behavioral and systemic physiological features mark preclinical Alzheimer’s disease

**DOI:** 10.64898/2026.10.26.761827

**Authors:** Liat Tsoran, Yuval Rom, Ahlam Taha, Yara Shwaiky, Fabienne Attal, Noa Naveh, Yoav Levy, Miriam Adam, Nina Fainstein, Naomi Habib, Tamir Ben-Hur

**Author notes:** These authors contributed equally (Liat Tsoran, Yuval Rom). These authors jointly supervised this work (Naomi Habib, Tamir Ben-Hur).

## Abstract

Alzheimer’s disease (AD) pathology accumulates years before overt cognitive impairment, yet its multidomain phenotype remains poorly defined. To limit confounding by uncontrolled comorbidities, we assembled a cohort of 644 older adults self-reporting intact cognition, free of systemic or neurological diseases and managed medically for risk factors. Plasma p-tau217 was elevated in 20%, and 93 of these were cognitively unimpaired. This biomarker-defined, risk factor - controlled preclinical AD subgroup showed subtle, concurrent cognitive and behavioral differences, accompanied by modest differences in physiological measures within clinical reference ranges, and alongside higher biomarkers of astrogliosis (GFAP) and neurodegeneration (NfL). Reported modifiable risk factor histories were not associated with p-tau217 when effectively managed. A proof-of-concept AI classifier integrating routine multidomain clinical data improved identification of p-tau217-high individuals relative to cognitive screening alone. These findings indicate preclinical AD is not phenotypically silent at the group level and support multidomain-based screening for biomarker testing and early-detection.

## INTRODUCTION

Late-onset Alzheimer’s disease (AD) is a progressive neurodegenerative disorder that develops over many years, with a prolonged preclinical phase that precedes overt cognitive impairment^1,2^. During this interval, amyloid-β accumulation is followed by tau pathology and synaptic dysfunction leading to progressive network disintegration that ultimately culminates in dementia^3^.

Yet, despite this long biological prodrome, AD is still commonly recognized only after the onset of cognitive decline, when substantial and largely irreversible neurodegeneration has already occurred^4^. Defining the earliest detectable features of AD before irreversible decline is therefore a central challenge for the development of effective disease-modifying and preventive therapies, risk stratification and timely intervention.

The emergence of blood-based biomarkers has transformed the ability to detect AD pathology prior to clinical symptoms. Plasma phosphorylated tau217 (p-tau217), is a sensitive and specific early biomarker of AD brain pathology that strongly predicts future cognitive decline^5–7^, Neurofilament light chain (NfL) is a biomarker of neurodegeneration and axonal injury, and glial fibrillary acidic protein (GFAP), is an astroglial activation biomarker. Together these three biomarkers provide a minimally invasive framework for interrogating ongoing pathological processes in the aging brain^8^. Specifically, they create an opportunity for identification of individuals with preclinical AD and test if there are any early manifestations of disease before functional decline becomes clinically apparent based on conventional diagnostic thresholds.

Resolving this question is challenging in typical aging cohorts, as aging populations frequently harbor multiple comorbidities, including vascular disease, metabolic dysfunction, systemic inflammation, that can independently influence cognition, behavior, and systemic physiology^9^. Brain co-pathologies may affect not only the timing of cognitive decline, but also its clinical and behavioral features^10^. As a result, subtle changes previously noted in preclinical AD or MCI^11^, may instead reflect common aging-associated illness and mixed neuropathologies. Conversely, early AD-associated signatures may be obscured by the broader biological heterogeneity of aging. Carefully selected, medically optimized and deeply characterized cohorts provide a complementary strategy for studying early AD by minimizing major confounding from uncontrolled comorbidities and enabling more precise interrogation of AD-associated cognitive, behavioral, and systemic changes.

Such cohorts are also important for clarifying the relationship between AD pathology and modifiable vascular and metabolic risk factors. Hypertension, diabetes mellitus, dyslipidemia, smoking, sleep disorders, and systemic inflammation were strongly associated with dementia risk^12^, but it remains unclear whether their association with early AD persists when these conditions are effectively monitored and treated. Most studies and meta-analyses reported very high prevalence of AD risk factors^13–16^, but either lack information on their actual control in the studied populations, or describe cohorts that are poorly controlled in regard to modifiable risk factors. Distinguishing reported risk-factor history from objectively uncontrolled physiological states may therefore help determine whether medical optimization attenuates the association between classical risk exposures and early AD pathology.

In parallel, growing evidence suggests that AD may not be confined to the brain, but may unfold within broader organism-wide physiological states involving metabolism, immune function, vascular health, and protein clearance^17^. However, whether such systemic physiological alterations are detectable at the preclinical stage, especially in individuals without overt systemic disease, remains unresolved. Identifying subtle systemic features associated with elevated AD biomarkers could provide insight into the broader biological context in which early AD develops and may also improve scalable approaches for identifying individuals who should undergo confirmatory biomarker testing.

Here, we assembled a medically optimized cohort of 644 adults aged 65–80 years who are self-reportedly cognitively healthy, functionally independent, and free of major neurological, psychiatric, or debilitating systemic disease. Moreover, they were medically monitored and treated for vascular and metabolic risk factors. Participants underwent detailed cognitive and behavioral assessment, medical and physiological profiling by blood tests, and measurement of plasma biomarkers p-tau217, NfL, and GFAP. We defined the subgroup of biomarker defined preclinical AD, as cognitively unimpaired participants with elevated p-tau217, and found that they associated with subtle cognitive, behavioral, and systemic physiological differences, alongside higher markers of neurodegeneration (Nfl) and astroglial activation (GFAP). However, at the individual level, clinical assessment alone could not identify the underlying AD pathology, in the pre-clinical AD as well as in AD-MCI participants. Reported histories of effectively managed vascular and metabolic risk factors showed limited association with p-tau217, whereas uncontrolled physiological states showed biomarker-specific associations. Finally, integrating routine multidomain clinical and medical data using an AI model, improved the identification of individuals with elevated p-tau217 compared with cognitive screening alone. Together, our findings derived from this unique medically optimized cohort show that biomarker-defined preclinical AD is embedded within a broad cognitive, behavioral and systemic physiological state, which can be integrated to a scalable screen for individuals at high risk for AD and assist biomarker-guided early AD detection at the population level.

## RESULTS

### A medically optimized aging cohort enables multidimensional characterization of preclinical AD, based on plasma biomarkers

To distinguish early Alzheimer’s disease (AD)-associated changes from vascular, and other neurological and systemic confounders, we established a tightly controlled medically optimized aging cohort of 644 individuals aged 65–80 years. Participants were self-reportedly cognitively healthy and functionally independent, lacked overt neurological, psychiatric, cerebrovascular, cardiovascular, renal, hepatic, malignant, or other major debilitating systemic diseases, and reported continuous monitoring and treatment of modifiable vascular and metabolic risk factors (**Fig. 1a–b**).

**Figure 1:**
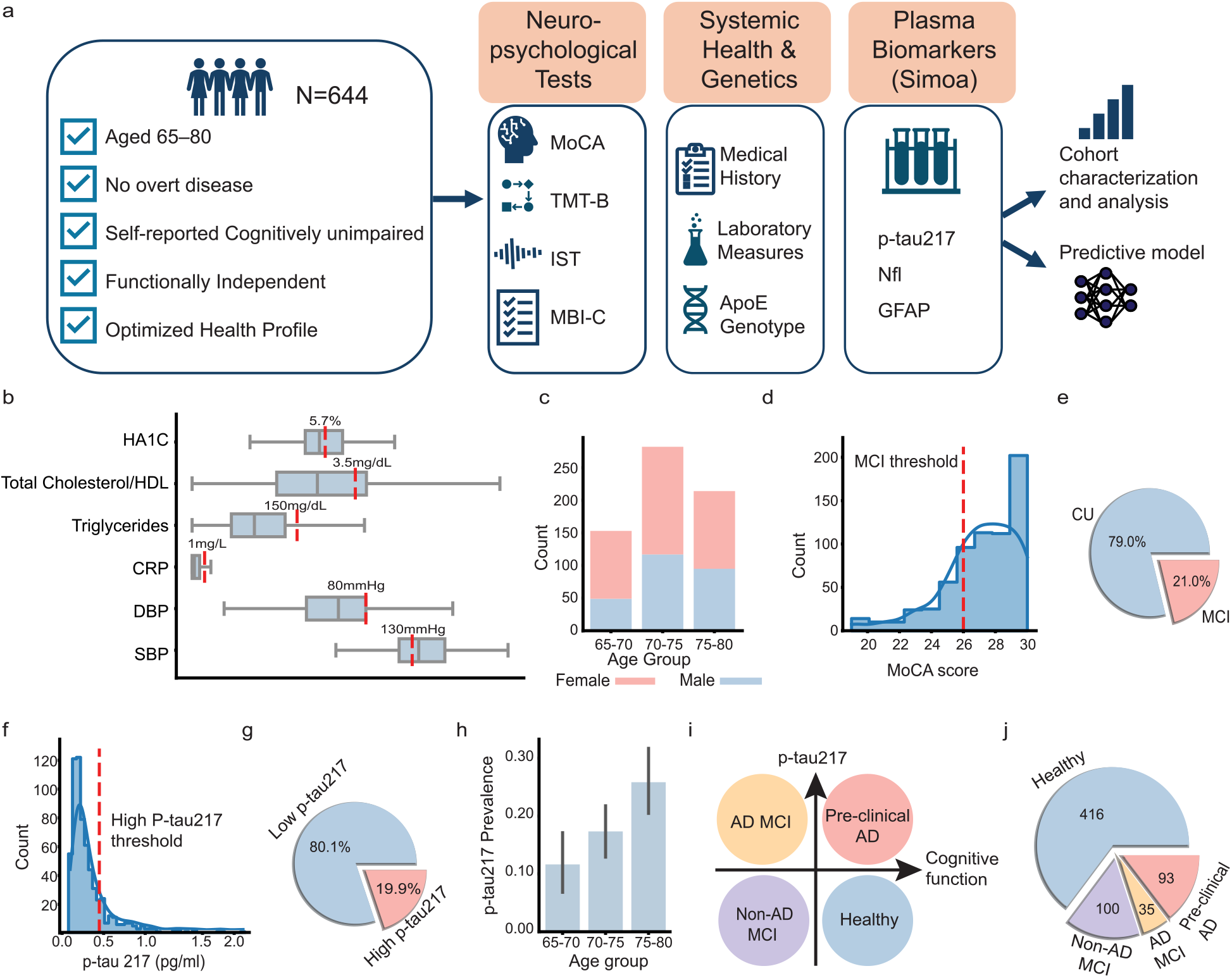
A medically optimized aging healthy cohort for studying preclinical AD based on plasma biomarkers, cognitive-behavioral tests, and clinical measurements. **(a)** Study design and inclusion criteria for the n=644 older adults participating who self-reported being cognitively unimpaired, functionally independent, and free of major diseases. **(b)** Box plots of baseline health parameters demonstrating the medically optimized state of the cohort. Red dotted line represents the clinical health guideline. Box spans 25^th^–75^th^ percentiles, the center line indicates the median, whiskers extend to 1.5 times the interquartile range. **(c)** Stacked bar plot showing the number of participants per age group and sex. **(d)** Distribution of Montreal Cognitive Assessment (MoCA) scores and MCI threshold (< 26, red line). **(e)** Percent of participants classified as cognitive unimpaired (CU) or MCI, with 21% falling below the MCI threshold. **(f)** Distribution of plasma p-tau217 levels and threshold (p-tau217 > 0.444 pg/mL, red line). **(g)** Percent of participants classified as high pTau-217, identifying 19.9% of the cohort with elevated AD-pathology markers. **(h)** Prevalence of elevated p-tau217 stratified by age group. **(i-j)** Stratification scheme (i) and distribution (j) of the cohort into healthy (n=416), preclinical AD (n=93), AD-MCI (n=35), and Non-AD MCI(n=100), based on the combined cognitive and biomarker thresholds.

Participants underwent comprehensive multidimensional clinical characterization, including structured medical history, detailed cognitive testing across multiple domains using the MoCA test^18^, Trail making test (TMT)-B^19^ and word fluency test^20^, as well as standardized behavioral assessment using the Mild behavioral Impairment (MBI) Checklist^21^, and functional evaluation by the Activities of Daily Living Inventory (iADL)^22^ questionnaire. Objective physiological profiling included routine laboratory measurements, cardiovascular parameters, and genetic testing for ApoE genotype for AD risk evaluation. Plasma biomarkers were quantified using Simoa technology for p-tau217, Nfl, and GFAP, enabling assessment of AD pathology, neurodegeneration, and astroglial activation in relation to cognitive, behavioral, and systemic health profiles.

Initial characterization confirmed that the cohort was functionally independent (median iADL=0) and medically well controlled for major risk factors, with most routine blood measurements within clinically accepted ranges (**Fig. 1b**, **Extended Data Table 1**). The cohort showed a balanced representation of men and women (**Fig. 1c**). Despite preserved self-reported cognitive function and functional independence, 21% of participants met MoCA-based criteria for mild cognitive impairment (MCI, MoCA<26), underscoring the high prevalence of unrecognized cognitive decline among the older population (**Fig. 1d-e**).

Plasma p-tau217 established cutoff (0.444 pg/ml^23^) classified 19.9% of the participants as positive for AD pathology (**Fig. 1f-g**). The prevalence of elevated p-tau217 increased with age, rising 1.49-fold per five-year interval (**Fig. 1h**). A more stringent p-tau217 cutoff (0.53 pg/ml^23^), yielded a similar 19.5% biomarker-high subgroup, supporting the robustness of the biomarker classification.

We next combined the cognitive and biomarker criteria to stratify participants into four groups: healthy (low p-tau217, cognitively unimpaired MoCA ≥26; n=416 participants), non-AD MCI (low p-tau217, MoCA<26; n=100), AD-MCI (high p-tau217, MoCA<26; n=35), and preclinical AD (high p-tau217, MoCA ≥26; n=93). This classification identified 93 preclinical AD individuals corresponding to 14.3% of the overall cohort, and 18.3% of the cognitively unimpaired participants (**Fig. 1i-j**). Of note, the prevalence of ApoE4 carriers in this new cohort was not higher than previously reported in other populations^24^, excluding the enrichment for APOE4 as the explanation for the high prevalence of preclinical AD.

Thus, despite self-reported cognitive health, functional independence, and medical optimization, a substantial subgroup showed biomarker evidence of AD pathology. This cohort therefore allowed us to test whether elevated p-tau217 is phenotypically silent before overt impairment or is already accompanied by detectable changes across biological, cognitive, behavioral, and systemic domains.

### Coordinated core pathology, neurodegeneration and astroglial activation biomarkers are detectable at the preclinical stage of AD

We next tested the relationships between plasma p-tau217, NfL and GFAP biomarkers, as supportive evidence for active biological processes in the brain at the preclinical stage, specifically indications for active neurodegeneration (NfL) and astroglial activations (GFAP) (**Fig. 2 a-b**). Consistent with previous studies, plasma levels of all three biomarkers were significantly correlated across the cohort (including NfL and GFAP: spearman R=0.53; p-tau217 and GFAP: R=0.42; **Fig. 2c-e**). Both p-tau217 and NfL were also significantly associated with lower cognitive performance across the cohort (**Fig. 2c**), suggesting that early cognitive changes in AD represent early neurodegenerative changes. We tested whether these biomarker associations were already present among cognitively unimpaired individuals. We found that elevated p-tau217 was associated with increased levels of GFAP and NfL (**Fig. 2f-g**), indicating that AD-pathology is accompanied by astroglial activation and neurodegenerative signals already at the preclinical stage^25–27^ that can be captured using plasma biomarkers.

**Figure 2:**
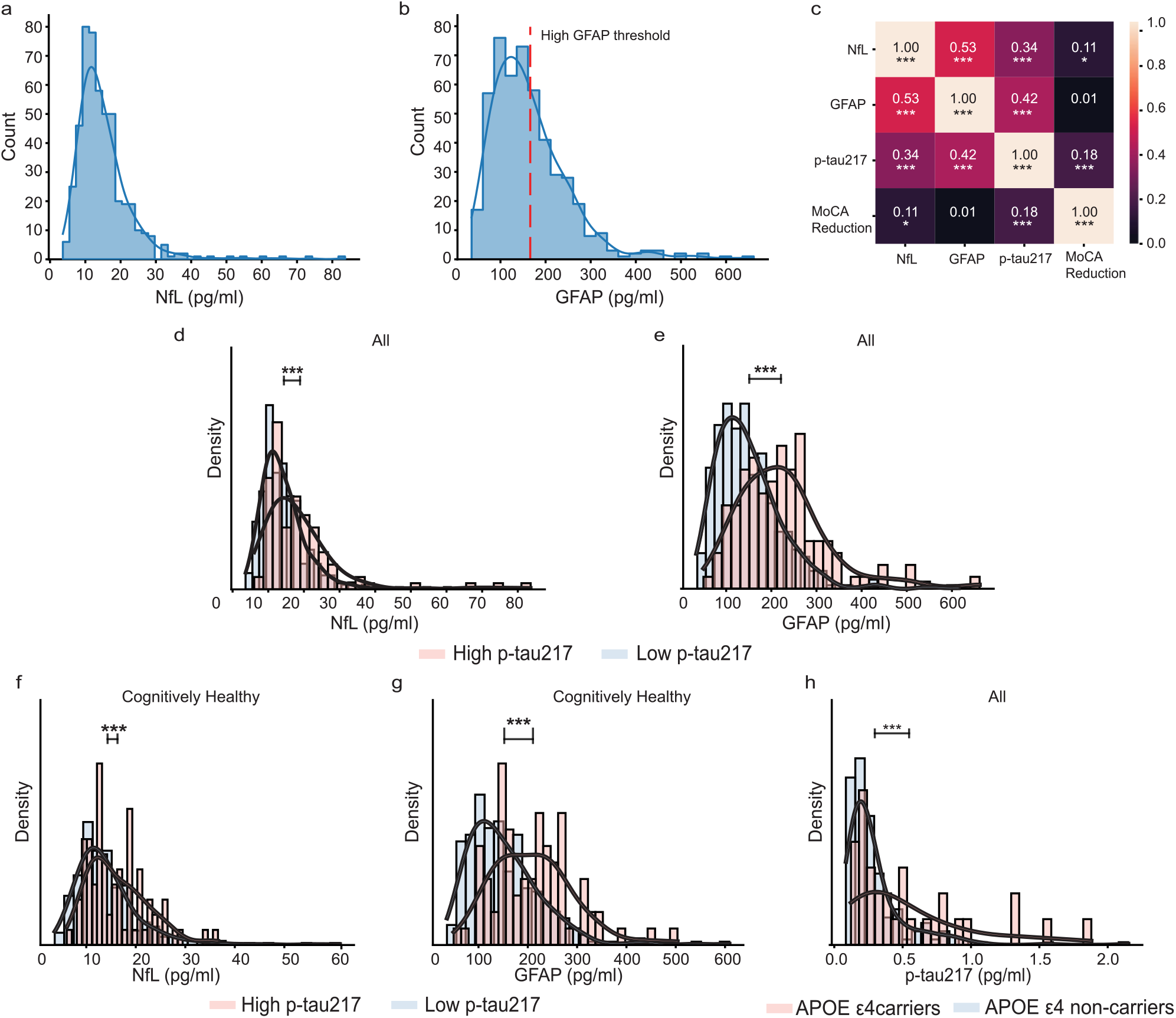
Plasma biomarkers associated with neurodegeneration and astroglial activation in preclinical AD. (a-b) Distribution of plasma neurofilament light chain (NfL, a) and glial fibrillary acidic protein (GFAP, b, dotted red line=166 pg/mL) concentrations across the cohort (n=510). **(c)** Biomarker correlations. Spearman correlations coefficients between MoCA score, plasma p-tau217, GFAP, and NfL; asterisks indicate significance levels. *<0.05; **<0.01; ***<0.005; **(d-e)** Density distributions of neurodegeneration marker NfL and astroglial marker GFAP stratified by p-tau217 status, showing elevated biomarker levels in participants with increased p-tau217 (n=510). **(f-g)** Density distributions of NfL and GFAP according to p-tau217 status within cognitively healthy participants (MoCA≥26, n=403), showing that these associations persist in the absence of overt cognitive impairment. **(h)** Plasma p-tau217 levels stratified by APOE ε4 carrier status, demonstrating significantly higher p-tau217 levels across the cohort among APOE ε4 carriers (Wilcoxon test).

Notably, carriage of high genetic risk allele ApoE4 was associated with elevated p-tau217, consistent with established increased AD risk, but was not associated with levels of neurodegeneration (NfL) or gliosis (GFAP) at this stage (**Fig. 2h, Extended Data Fig. 1a-b**).

Together, these findings show that elevated p-tau217, indicating accumulating AD pathology, is not an isolated event, but is accompanied by elevated biomarkers of astroglial activation and neurodegeneration, further supporting a distinct biological state in the brain even among cognitively unimpaired individuals. We therefore next asked whether this biological state is also associated with subtle cognitive and behavioral manifestations that remain below conventional clinical thresholds.

### Concurrent cognitive and behavioral alterations accompany elevated p-tau217 before overt clinical impairment

We next examined whether individuals with preclinical AD already exhibit detectable cognitive and behavioral alterations (**Fig. 3a-f**). Across the cohort, higher p-tau217 levels were associated with lower cognitive performance scores across tests (**Fig. 3a**). Consistently, preclinical AD participants demonstrated subtle but significant reductions in global cognitive performance, as compared with control (biomarker-low, cognitively normal) participants, despite MoCA scores remaining within the clinically normal range (**Fig. 3b**). Domain level analysis of the cognitive tests indicated that these differences were characterized by slower executive processing (Trail Making Test–B, TMT-B), reduced semantic fluency (IST), and selective impairment in frontal-type memory retrieval and global memory, while other domains were relatively preserved (**Fig. 3b-c**).

**Figure 3:**
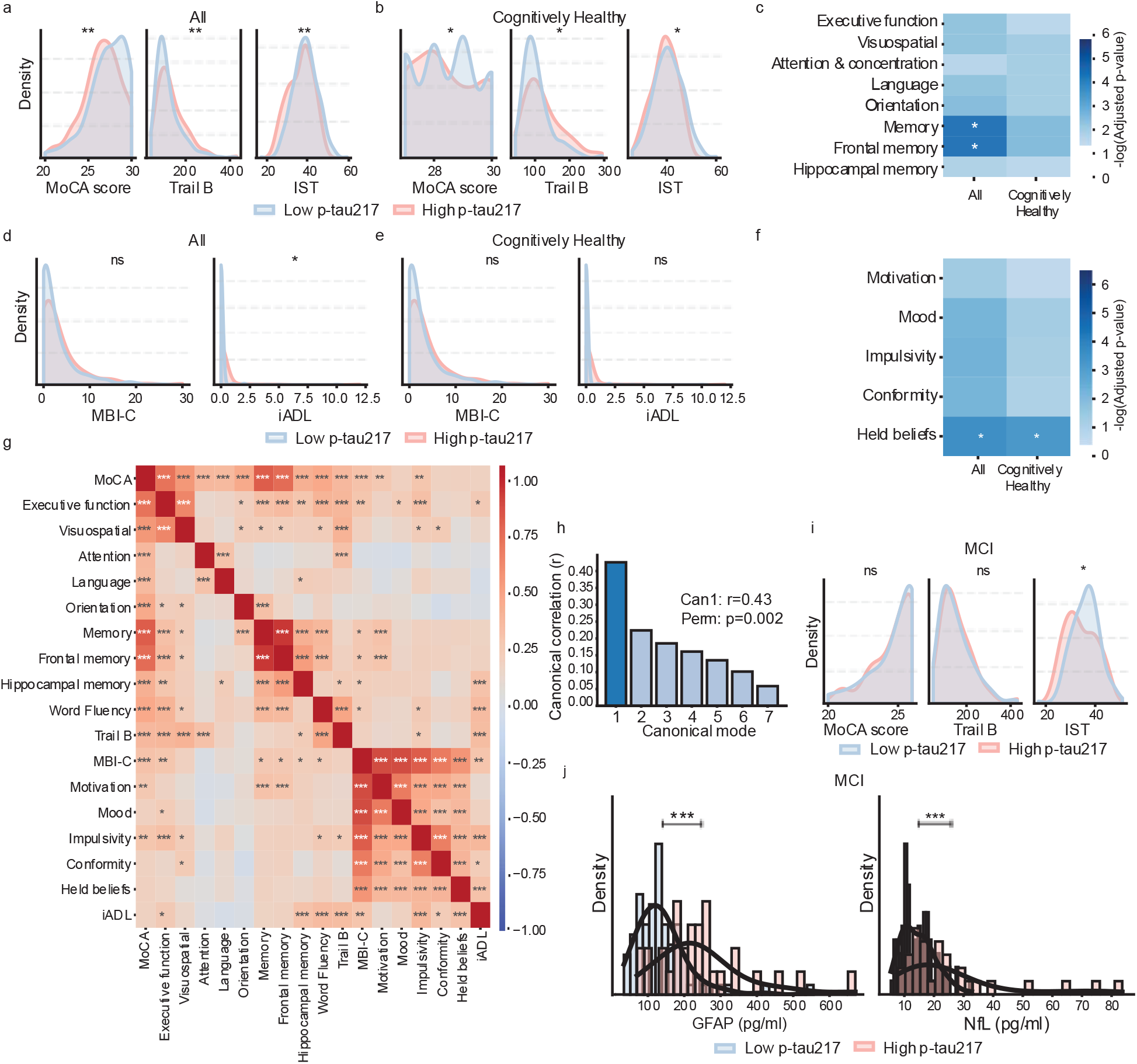
Concurrent cognitive and behavioral alterations emerge at the preclinical stage of Alzheimer’s disease. **(a-b)** Distributions of cognitive performance measures (MoCA, TMT-B, and IST) stratified by p-tau217 status in the full cohort **(a)** and cognitively healthy participants **(b).** High p-tau217 was associated with poorer performance in all three tests. **(c)** Heatmap showing the -log_10_ FDR adjusted P-values Mann Whitney U test for group (high-vs. low-p-tau217) comparisons across cognitive domains, highlighting selective impairments in executive function and frontal-type memory retrieval in preclinical AD. (**d-e)** Distributions of behavioral measures (MBI-C score and iADL) stratified by p-tau217 status in the full cohort **(d)** and cognitively healthy participants **(e)**. **(f)** Heatmap showing the negative -log_10_ FDR adjusted P-values Mann Whitney U for group (high-vs. low-p-tau217) comparisons across behavioral domains, including: interest, motivation, and drive (Motivation); mood or anxiety symptoms (Mood); ability to delay gratification and control behavior, impulses, oral intake and/or changes in reward (Impulsivity); following societal norms and having social graces, tact, and empathy (Conformity); strongly held beliefs and sensory experiences (Held beliefs). Highlighting increased endorsement of suspiciousness and unrealistic beliefs in high p-tau217 individuals. (**g**) Pearson correlation matrix depicting associations between behavioral indices and cognitive scores within the high p-tau217 subgroup. Note that multiplied the cognitive parameters (MoCA, Executive function, Visuospatial, Attention, Language, Orientation, Memory, Frontal memory, Hippocampal memory, Word Fluency) by negative 1, thus positive correlations reflect a reduction in cognitive performance. (**h**) Canonical correlation analysis (CCA) suggests co-occurrence of cognitive and behavioral variables within the cohort. The first canonical vector (can1) reached 0.43 spearman rho correlation for cognitive measures, higher loadings correspond to lower cognitive performance, permutation test (perm) was significant with P-value=0.02. **(i)** Distributions of cognitive performance measures (MoCA, TMT-B, IST) within participants meeting criteria for MCI (MoCA score <26), stratified by p-tau217 status. Standard clinical assessments (MoCA, TMT-B) did not differ significantly between AD-related MCI from non-AD MCI, except for mildly worse performance in word fluency (IST) in MCI with high p-tau217 levels. **(j)** Distributions of plasma NfL and GFAP levels within the MCI subgroup, stratified by p-tau217 status. Plasma GFAP and NfL level were higher in AD-MCI than in non-AD MCI. For all panels: *<0.05; **<0.01; ***<0.005.

Behavioral alterations paralleled these cognitive changes. Across the entire cohort, high p-tau217 was associated with higher Mild Behavioral Impairment Checklist (MBI-C) scores and with higher incidence of iADL>0 (**Fig. 3d)**. Although total MBI-C scores did not differ significantly between preclinical AD and healthy individuals (**Fig. 3e**), domain-level analysis revealed a selective increase within preclinical AD in strongly held beliefs and sensory experiences, linked to increased endorsement of suspiciousness and unrealistic beliefs (**Fig. 3f**). These subtle behavioral changes might infer early frontal lobe dysfunction, indicated by increased dogmatic thinking, mental rigidity, and mildly decreased reality monitoring and ability to interpret others’ intentions^28^. Notably, within the biomarker-positive (high p-tau217) subgroup, cognitive performance and behavioral scores were significantly correlated (p<0.001; MBI-C and MoCA, Pearson R = 0.26; strongly held beliefs and Trail-B, R = 0.31; **Fig. 3g**). Canonical correlation analysis further showed that both cognitive and behavioral measures loading, loaded onto a shared canonical mode of variation persisted robustly across the entire cohort (**Fig. 3h, Extended Data Fig. 2a-d**). Thus, these behavioral changes are concurrent with early cognitive decline, already at the preclinical stage of AD, rather than forming a fully independent trajectory. Moreover, the specific tests associated with preclinical AD in both the cognitive and behavioral domains, including poorer executive functions and suspicious and unrealistic beliefs, may be both linked to subtle frontal lobe dysfunction.

Together, these findings challenge the view that preclinical AD is clinically silent, indicate that elevated p-tau217 is accompanied by subtle cognitive and selected behavioral differences before overt cognitive impairment at the group level, which might reflect coordinated manifestations of the same underlying pathological process. These changes were detectable at the group level but remained modest, and cannot be the basis of reliable diagnosis, indicating an early multidomain phenotype rather than a clinically recognizable syndrome.

### Clinical assessment alone does not distinguish AD-related MCI from non-AD MCI

Although subtle cognitive and behavioral differences accompanied elevated p-tau217 before overt impairment, it remained unclear whether standard clinical evaluation could identify the etiological basis of cognitive impairment and the underlying pathology at the individual level. We compared individuals meeting criteria for MCI with elevated plasma p-tau217, defined as AD-MCI, versus those with MCI but low p-tau217 levels, defined as non-AD-MCI. Standard cognitive and behavioral assessments failed to distinguish these groups (**Fig. 3i, Extended data Fig. 2e**). AD-MCI and non-AD-MCI participants showed comparable performance, including global MoCA scores, TMT-B and MBI-C behavioral assessment scores and MoCA test domain-level profiles and subtle difference in IST scores (**Fig. 3i**, **Extended Data Fig. 2e, Source data statistics**). In contrast, plasma biomarkers (GFAP, and NfL) clearly separated AD-MCI from non-AD-MCI (p<0.001; **Fig. 3j**). AD-MCI participants, showed higher GFAP and NfL levels relative to non-AD MCI participants (p<0.001; **Fig. 3j**), indicating that AD pathology, marked by p-tau217, is accompanied by higher levels of astroglial activation and neurodegeneration. Conversely, the lower NfL levels observed in the non-AD MCI suggest that reduced cognitive performance in this group may arise from distinct mechanisms and, in many participants, may not reflect active neurodegeneration.

Thus, early AD-related differences are detectable at the group level, but cognitive and behavioral assessment alone did not determine the etiology of clinically measurable impairment. This distinction underscores a critical limitation of symptom-based diagnosis at early disease stages and the complementary roles of multidomain phenotyping and blood biomarkers: clinical features can reveal early change, whereas blood-based biomarkers are required for etiological classification and identification of the underlying AD pathology.

### Current metabolic control, rather than risk-factor history, associates with AD-related biomarkers

The presence of elevated p-tau217 in otherwise medically optimized individuals next prompted us to examine whether conventional vascular and metabolic risk factors remained associated with AD-related biomarkers when their current physiological control was considered (**Supplementary Table 1**). Surprisingly, neither the presence nor the cumulative number of reported risk factors was associated with plasma p-tau217 levels. The prevalence of common risk factors, including hypertension, diabetes mellitus, hyperlipidemia, ischemic heart disease, and sleep disorders, was comparable between individuals with elevated and low p-tau217 levels, as well as between preclinical-AD and healthy individuals (**Fig. 4a**). Smoking was the exception, both any smoking history and heavy smokers, defined as current smoking with >10 pack-years, were associated with elevated p-tau217 levels across the entire cohort and in preclinical AD compared to healthy controls (**Fig. 4a**).

**Figure 4:**
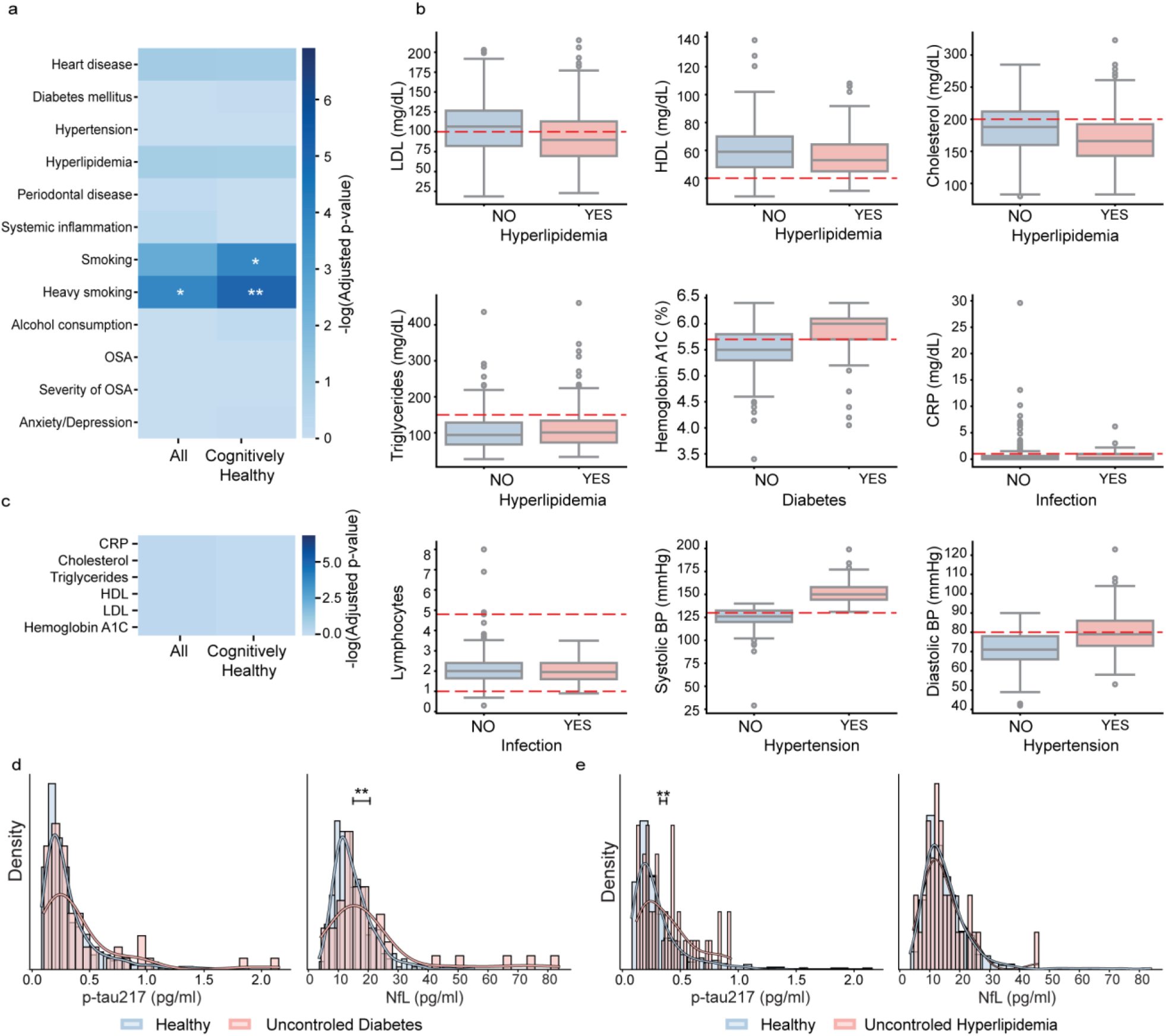
Controlled modifiable risk factors show limited association with biomarker-defined AD pathology. **(a)** Heatmap showing the negative -log_10_ adjusted P values for associations between self-reported modifiable risk factors and high pTau-217 status in the full cohort and preclinical AD status in the cognitively unimpaired subgroup. Most risk factors showed no significant association with elevated p-tau217, with only smoking and heavy smoking (above 10 pack years) reaching significance. Asterisks denote adjusted P < 0.05. **(b)** Box plots of objective laboratory measurements (LDL, HDL, cholesterol, triglycerides, HbA1C, CRP, lymphocytes, SBP, and DBP) stratified by self-reported disease status, demonstrating generally well-controlled cardiovascular and metabolic risk factors among participants reporting hyperlipidemia, diabetes, infection, or hypertension. Box spans 25th–75th percentiles, the center line indicates the median, whiskers extend to 1.5 times the interquartile range. **(c)** Heatmap showing the negative -log_10_ adjusted P values for associations between laboratory measurements and high pTau-217 status. **(d)** Density distributions of plasma p-tau217 and NfL according to diabetes control status. Uncontrolled diabetes (HbA1c>6.5%) was associated with elevated NfL levels but not with p-tau217. **(e)** Density distributions of plasma p-tau217, and NfL according to hyperlipidemia control (LDL>116 mg/dL, non-HDL cholesterol > 146 mg/dL). Uncontrolled hyperlipidemia was associated with elevated p-tau217 levels but not with NfL.

Due to the lack of association between reported risk factor history and AD biomarkers, we examined objective evidence of medical control of the risk condition by laboratory and physiological measurements. As we expected within our cohort, the majority of participants reporting vascular or metabolic risks history were medically well controlled (**Fig. 4b**): Participants reporting diabetes generally had hemoglobin A1C levels within or near recommended treatment targets; those reporting hyperlipidemia history had total cholesterol and LDL levels within guideline ranges, and often lower than participants without reported hyperlipidemia, consistent with effective pharmacological management; and participants reporting hypertension showed controlled systolic and diastolic blood pressure values (**Fig. 4b-c**). These findings indicate that self-reported risk-factor history alone, without updated physiological status, may not capture the physiological links relevant to early AD biology.

We therefore stratified participants by measurable risk status, distinguishing medically controlled from uncontrolled states based on the relevant blood measurements. Because most risk factors were effectively controlled in our cohort, only a limited number of participants met criteria for uncontrolled disease. Nevertheless, we could show biomarker-specific associations for diabetes mellitus and hyperlipidemia risk history, when blood profiles indicated measurable dysregulation. Participants with abnormal hemoglobin A1C levels (HBA1C>6.5%) exhibited elevated plasma Nfl, whereas participants with controlled diabetes did not (**Fig. 4d**); this association was not observed for plasma p-tau217 or GFAP (**Fig. 4d, Extended Data Fig. 3a**). Conversely, though self-reported hyperlipidemia was not associated with p-tau217 levels (**Fig. 4a**), uncontrolled dyslipidemia, defined by guideline-based LDL and non-HDL cholesterol thresholds (LDL >116 mg/dL and non-HDL cholesterol >146 mg/dL), was significantly associated with higher p-tau217, but not with GFAP or NfL (**Fig. 4e, Extended Data Fig. 3b**). These patterns position each of these risk factor along different axes of the disease cascade, suggesting uncontrolled diabetes mellitus associate with neuronal damage, independent of AD-pathology^29,30^, while uncontrolled hyperlipidemia is linked to AD pathology^31^.

Together, these findings suggest that reported risk-factor history alone incompletely captures the systemic conditions associated with early AD-related biomarkers. In this medically optimized cohort, most treated risk-factor histories showed limited association with p-tau217, whereas contemporaneous evidence of metabolic dysregulation from blood tests showed biomarker-specific associations.

### Subtle systemic physiological signatures accompany preclinical AD

Because conventional risk-factor diagnoses showed limited association with elevated p-tau217 when medically controlled, we next asked whether subtle variations in routine physiological measures could capture a broader systemic context associated with early AD pathology (**Extended data Table 1**). Since individuals with overt systemic disease were excluded from our cohort and most laboratory measurements fell within clinically accepted reference ranges (**Fig. 1b**, **Extended Data Table 1**), our cohort enabled assessment of subtle physiological differences associated with preclinical AD in otherwise medically optimized individuals. Individuals with elevated p-tau217 showed modest but consistent systemic physiological differences. Plasma creatinine levels were slightly higher in p-tau217-high individuals, with corresponding trends in estimated glomerular filtration rate (GFR) (**Fig. 5a**), although values remained within clinically normal limits and participants with overt renal failure were excluded from the study. Thus, these differences reflect variations within the physiological range rather than clinically significant renal impairment. Body mass index (BMI) was also slightly lower in the preclinical AD group, an effect that was more pronounced in men (**Fig. 5b**). Because BMI values remained within the non-cachectic range, and the shifts were small in magnitude, this is consistent with subtle systemic metabolic modulation rather than overt weight loss.

**Figure 5:**
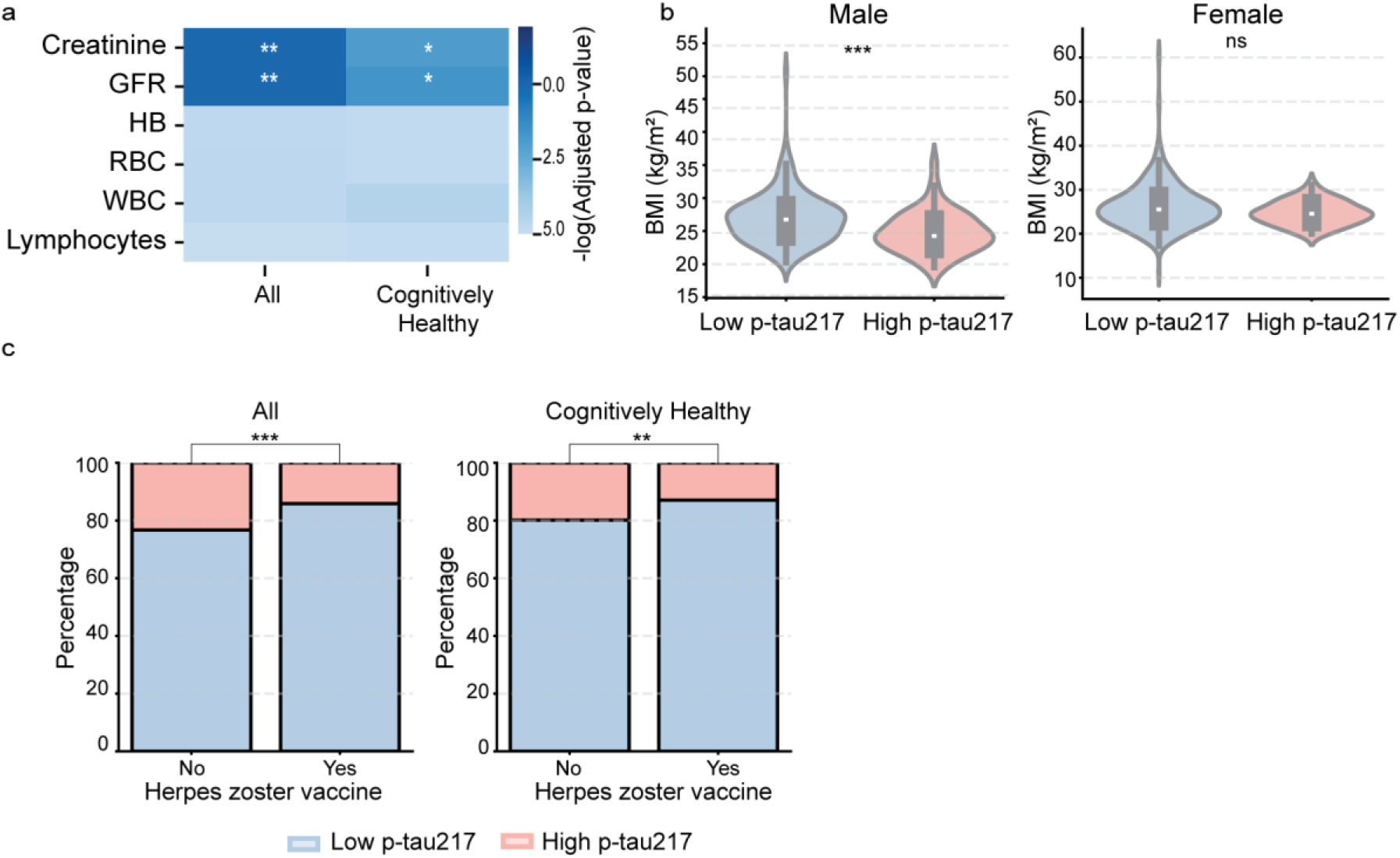
Subtle systemic physiological signatures accompany preclinical AD. **(a)** Heatmap showing the -log_10_ FDR adjusted P-values Mann Whitney U test, modest variations within clinically normal ranges for renal measures (Creatinine, GFR) in participants with high p-tau217, and cognitively healthy individuals with high p-tau217. **(b)** Body mass index (BMI) distributions stratified by sex, showing lower BMI among participants with high p-Tau217, particularly in males. **(c)** Bar charts illustrating significantly lower frequency of prior herpes zoster vaccination among individuals with elevated p-tau217 compared to the low p-tau217 baseline.

Finally, prior herpes zoster vaccination was less frequent among individuals with elevated p-tau217 (**Fig 5c**), consistent with emerging epidemiological evidence linking zoster vaccination history to reduced dementia risk^32^. Although observational, this association raises the possibility that immune history or vaccination-related protection may relate to early AD risk, beyond the prior link to clinical symptoms of dementia.

Thus, elevated p-tau217 was accompanied by subtle systemic physiological differences extending beyond cognitive and behavioral measures. Although these features were individually modest, largely within clinical reference ranges, and not necessarily specific to AD, their convergence across domains suggested a broader systemic context for early AD, and indicated they might become informative when considered jointly.

### An integrative model of routine multidomain clinical data enriches identification of elevated p-tau217

The preceding analyses identified modest differences at the group level across cognitive, behavioral, medical, and physiological domains, none of which was individually sufficient to identify elevated p-tau217 at the individual level. We therefore tested whether integrating these distributed signals could improve the prioritization of individuals for biomarker testing. We constructed an integrative prediction model, incorporating demographic variables, cognitive and behavioral measures, reported risk factors, and systemic routine laboratory parameters (**Fig. 6a**). As a clinical-based reference, we classified participants according to standard global cognitive performance, using a slightly stricter threshold of MoCA≤26 (including MCI and individuals with borderline cognitive performance). This cognitive only approach showed limited results, identifying 45% of individuals with elevated p-tau217, missing the remaining 55% who had MoCA scores≥27, and with high rate of false positives (30%, **Fig. 6b**). This aligns with our finding that around 70% of MCI individuals display low p-tau217 (**Fig. 1j**). Thus, global cognitive screening alone showed limited sensitivity to discriminate underlying AD pathology, particularly limiting its utility in cognitively unimpaired individuals.

**Figure 6:**
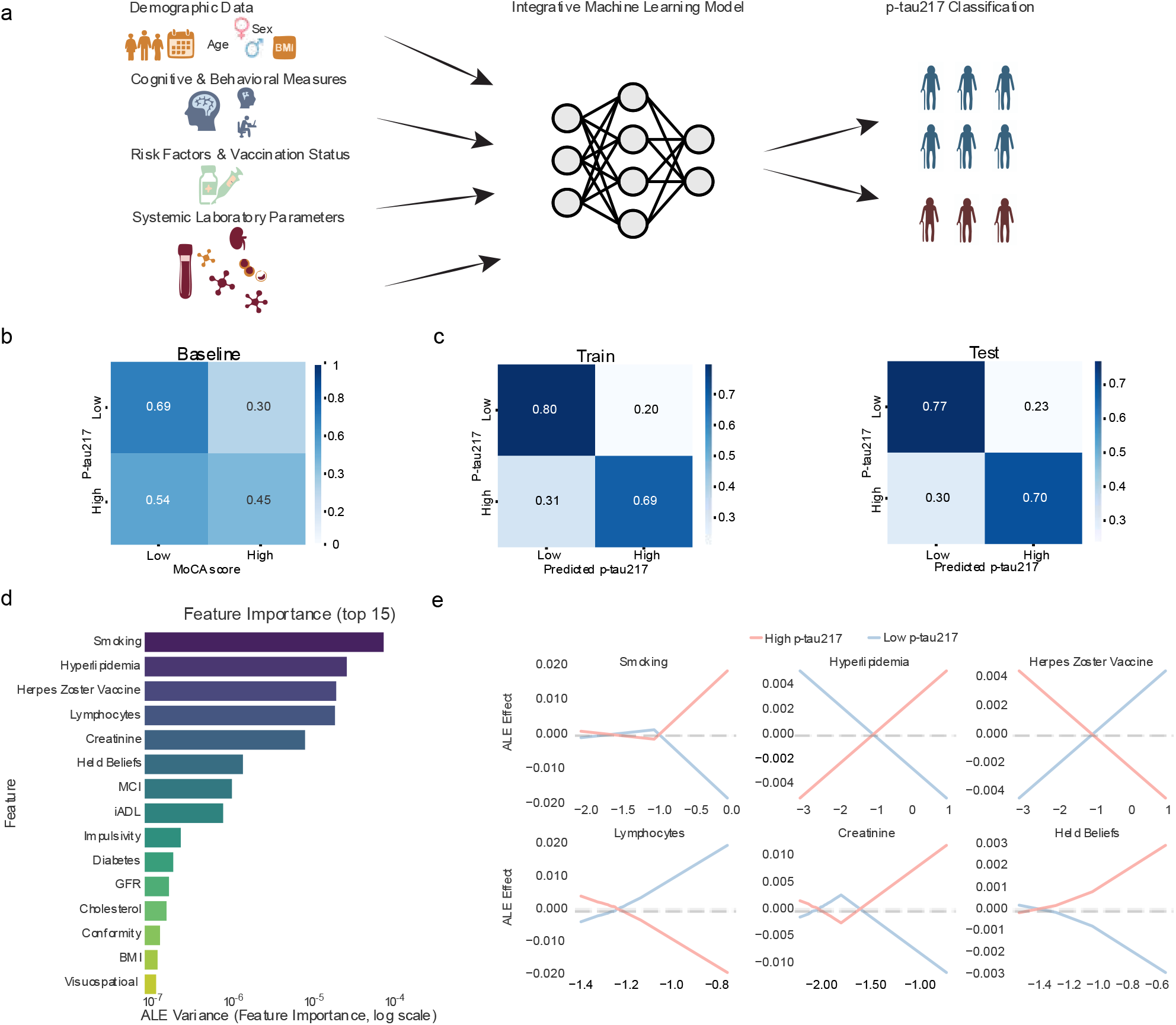
An integrative model predicts individuals with plasma biomarker defined AD pathology from routine clinical data. **(a)** Schematic overview of the integrative prediction framework. Demographic variables, cognitive and behavioral measures, risk factors and vaccination status, and systemic laboratory parameters were used to classify participants according to plasma p-tau217 status. Comparison of test set recall (true positive rate) between the MoCA-based baseline classifier and the integrative model. Train-validation-test split 70:15:15. **(b)** Confusion matrix for the MoCA-based baseline classifier (MoCA=26), normalized by row. **(c)** Training and test confusion matrices for the TabPFN model (**methods**), normalized by row. **(d)** Feature importance ranking based on Accumulated Local Effects (ALE) variance, presented in log scale. **(e)** Accumulated Local Effects (ALE) profiles for the six highest-ranked features identified by the TabPFN model, illustrating the contribution of feature values to model predictions.

We next trained machine-learning models to classify participants as p-tau217-high or p-tau-low using the multimodal clinical dataset. Models were trained on 70% of participants and tested on the 30% held-out test set (**Fig. 6a**). Across the evaluated architectures, both the tabular foundation model TabPFN^33^ and logistic regression model significantly outperformed the cognitive-only baseline across AUC, recall, and F1 scoring metrics (**Fig. 6c**, **Extended Data Fig. 4a-d**). The positive predictive value increased from prevalence of 20% corresponding to the prevalence of p-tau217 high, to 27% using the standard MoCA-based classifier, and to 46% using our integrative model. The prediction true positive rate (recall) increased from 45% with the cognitive-only classifier to 70% with the integrative model. This improvement substantially reduced the number of confirmatory biomarker tests required to identify p-tau217-high individuals in a population screen. Based on the observed positive predictive values of the model and the observed prevalence of p-tau217, identifying 100 p-tau217-high individuals through *untargeted screening* would require approximately 500 participants, compared with approximately 143 tests among individuals prioritized by the *integrative model* - a 71% reduction in biomarker testing burden. Thus, integrating multiple weak clinical and physiological signals improved both the sensitivity and efficiency of p-tau217 identification relative to cognitive screening alone.

To determine which features contributed to the model predictions, we performed post-hoc feature-importance analysis on the trained TabPFN model using Accumulated Local Effects^34^ (ALE) analysis (**Fig.6d**, **Extended Data Fig. 4e**, **Methods**). Systemic physiological features consistently ranked among the strongest contributors and generally exceeded cognitive measures, including MCI-status, in predictive importance (**Fig. 6d-e**). Highly ranked features included hematologic parameters, renal and metabolic factors; smoking and herpes zoster vaccination status; selected behavioral measure, such as strongly held beliefs and impulsivity; MCI status and visuospatial functions (**Fig. 6d**). Logistic regression model provided an interpretable linear benchmark for informative features, and identified overlapping feature classes, including lymphocytes abundance, BMI, GFR, HbA1C, cholesterol, smoking, and several cognitive and systemic features.

Lymphocyte count illustrates how a feature may contribute to the multivariate classification despite showing only a modest group level difference. Although overall lymphocyte counts did not differ significantly by p-tau217 status (**Extended Data Fig. 4f**), clinically defined lymphopenia (low lymphocyte counts of <1×10⁹ /L) was more frequent at higher p-tau217 levels (P < 0.005; **Extended Data Fig. 4g**). This suggests that immune-related information weakly associated with p-tau217 may contribute to classification in combination with other multidomain features.

Overall, these findings provide a proof of concept that integrating weak signals distributed across routine cognitive, behavioral, and physiological can increase the sensitivity and precision of the identification of individuals with elevated p-tau217 beyond global cognitive screening alone. This approach is not intended to diagnose AD or replace plasma biomarkers, but may provide a scalable strategy for prioritizing individuals for confirmatory biomarker testing and early-intervention studies.

## DISCUSSION

Our findings identify a multidomain phenotype associated with biomarker-defined AD pathology before overt cognitive impairment. In functionally independent, medically monitored and risk factor-treated older adults, elevated plasma p-tau217 was accompanied by subtle differences in cognition, behavior, markers of astroglial activation and neurodegeneration, and systemic physiology. Yet these individual features were modest and did not provide etiological classification at the individual level, but when combined in a multidomain integrative model they could enrich for individuals with elevated pTau217. Moreover, risk-factor history alone showed limited association with AD biomarkers when effectively managed. Rather, association of modifiable vascular and metabolic risk factors with early AD depends strongly on their current physiological control. These results highlight the distinction, between a phenotype that is detectable at the group level and one that is clinically diagnostic, and supports the use of blood biomarkers to establish underlying AD pathology, while suggesting that integrated routine clinical data could help prioritize individuals for confirmatory biomarker testing.

Our study dealt with a central challenge in studies of older adults, commonly carrying comorbidities that can independently affect cognition, behavior, circulating biomarkers, and systemic physiology. By studying a medically optimized aging cohort, we sought to reduce this confounding and isolate early preclinical AD-related clinical and physiological changes. We found approximately 20% prevalence of AD pathology in this apparently healthy aging cohort, and approximately 15% met the criteria for preclinical AD. These findings underscore the high prevalence of AD brain pathology in the aging population, even in risk factor-controlled individuals, and support the need for scalable screening for early detection.

The cognitive and behavioral profile observed in preclinical AD showed subtle, coordinated, decline in specific cognitive and behavioral domains. These align with reports that sensitive neuropsychological measures can detect early deviations years before overt clinical impairment^35,36^, and extend them by showing, in a medically optimized cohort, that these changes are specifically associated with AD pathology rather than common comorbidities. Importantly, we now show that both cognitive and behavioral selective domains found to be affected at preclinical AD, such as frontal-type memory retrieval and increased suspicious and unrealistic beliefs, converging together to early Frontal lobe dysfunction^37^. This suggests that behavioral and cognitive impairments may not represent two independent prodromal syndromes, as previously suggested^38^, but rather coordinated manifestations of shared underlying pathology^39^.

At the same time, our findings highlight a critical limitation of symptom-based classification. Once cognitive impairment was clinically measurable by standard thresholds (*i.e.* the MCI individuals within our cohort), standard cognitive and behavioral assessments could not distinguish AD-MCI from non-AD-MCI^40^, whereas plasma biomarkers separated these groups^41^. The inability to differentiate clinically between AD and other causes of MCI is partly explained by the variability in clinical presentation of AD, related to the pattern of regional distributions of brain pathology^42^, affecting different cognitive domains. This timely finding emphasizes that biomarker-based classification is required for etiological diagnosis and appropriate targeting of prevention and early-intervention trials.

In this medically optimized cohort, histories of hypertension, diabetes, and hyperlipidemia were not associated with elevated p-tau217, contrary to their established link to AD risk^43^. While accumulating research indicate that vascular and metabolic risk management can lower the risk of dementia^12,44,45^, our results indicate that effective risk management may attenuate their association with AD pathology, even at the preclinical stage. Moreover, diagnostic history alone was insufficient to determine AD risk, while objective physiological measurements were more informative.

Our findings also associate risk and protective factors with distinct pathogenetic events. Although observational and limited by the small number of uncontrolled cases, our findings suggest that uncontrolled dyslipidemia may promote p-Tau217-associated brain pathology prior to cognitive decline^46,47^, whereas uncontrolled diabetes may promote Nfl-associated neurodegeneration independent of AD pathology^48,49^. Furthermore, the negative association between herpes zoster vaccination and elevated p-tau217 extends prior epidemiological links between zoster vaccination and reduced dementia risk to a biomarker-defined preclinical AD stage^32^. Although the causal protective effect of vaccination on preclinical AD requires further validation, this finding is consistent with the possibility that immune-training-related protection, or infection-related mechanisms may influence p-tau217-associated AD pathology upstream of clinical symptoms onset^50^.

Beyond classical risk factors, we identified modest but consistent systemic physiological changes associated with preclinical AD, including hematologic parameters, renal^51^, and body mass index^52–54^. These values largely remained within clinically normal ranges and therefore should not be interpreted as overt systemic disease. Specifically, renal differences were small and participants with renal failure were excluded, arguing against clinically significant impairment of renal functions as modulator of plasma biomarker levels^55,56^. These findings suggest that early AD pathology may emerge within a broader organismal context characterized by subtle systemic alterations, as previously shown for chronic inflammation and immune aging^57–59^. Whether such systemic features contribute to AD pathogenesis, arise as consequences of early brain pathology, or reflect shared upstream aging processes cannot be determined from the present cross-sectional data. These findings emphasize that AD may evolve within interconnected organism-wide physiological states that are individually subtle but become informative when integrated across domains.

Our integrative prediction model demonstrates the practical value of multidimensional routine clinical data. Combining routine blood measures, medical history, cognitive tests, behavioral questionnaires, and demographics predicted elevated pTau217, substantially better than cognitive testing alone. Our model improves and extends beyond previous classifiers trained to predict convergence to AD-dementia and not designed to predict existing AD pathology^60,61^. Within our models, systemic features contributed more than individual cognitive measures, further supporting a broader systemic context for preclinical. Although not intended to replace plasma biomarkers, this approach may provide a scalable screening funnel to prioritize confirmatory biomarker testing in population-level prevention programs.

Several limitations should be considered. First, the cross-sectional design precludes inference regarding the temporal order or causality, requiring longitudinal follow-up. Second, although the medically optimized cohort reduced comorbidity-related confounding, it may limit generalizability to more heterogeneous populations. Future longitudinal studies integrating biomarkers, imaging, and immune profiling may clarify links between systemic aging and neurodegeneration.

Together, our findings show that preclinical AD unfolds within a broader systemic context beyond the brain, marked by coordinated cognitive, behavioral, and systemic changes. Integrating multidimensional routine clinical and physiological data with plasma biomarkers may enable scalable early detection, risk stratification, and prevention-oriented intervention before irreversible neurodegeneration.

## METHODS

### Cohort recruitment and assembly

#### Inclusion and exclusion criteria

Inclusion criteria: 65-80 years old, self reportedly cognitively unimpaired and functional people and no history or current features of other neurodegenerative, cerebrovascular and neuropsychiatric diseases. Exclusion criteria: people with extrapyramidal features, focal signs or history of stroke, and overt affective and behavioral changes which may point at primary Frontal lobe syndromes. We also excluded people with major active systemic diseases, such as cancer, renal failure and severe heart failure

#### Method of recruitment

An open call inviting people aged 65-80 who are self-reportedly cognitively and functionally unimpaired to join this study was distributed via electronic media and by lectures to the public.

#### Demographic, medical, cognitive and behavioral evaluation

Demographic data included age, gender, years of education, family status, and country of origin. Medical data included measuring blood pressure and a questionnaire on existence of heart disease, diabetes mellitus, hypertension, hyperlipidemia, periodontal disease, recurrent infections (such as Pneumonia and urinary tract infections), systemic inflammatory diseases, smoking (current and past), alcohol consumption, insomnia, obstructive sleep apnea, physical activity, cognitive activity, dementia in the family, and vaccination for Herpes Zoster. Data on routine blood test results was obtained for: Hemoglobin and % Hemoglobin A1C, total Cholesterol, Triglycerides, LDL, HDL, white blood cell count, Lymphocyte count, Creatinine and C-reactive protein (CRP). Cognitive evaluation was performed using the MoCA test^18^, with MoCA≥26 considered normal, and a score of 19-25 considered as mild cognitive impairment (MCI). A few participants that obtained a MoCA score <19 were excluded from analyses. We implemented a variation on the common division of MoCA testing into cognitive domains of executive, spatial, attention and concentration, memory, language and orientation (**extended data Table 2**) In addition, we scored retrieval of the 5 memory items by cue or recognition from multiple choices to define Frontal type memory impairment versus Limbic type with no recognition (**Extended Data Table 2**). We also performed Trail making test (TMT)-B^19^, and word fluency using Isaac stress test^20^ given studies suggesting that these tests can reveal changes that precede up to a decade the clinical diagnosis of dementia^36^. Behavioral evaluation was performed using the Mild behavioral Impairment (MBI) Checklist and its breakdown to the various domains^21^. The MBI-C domains evaluated include: interest, motivation, and drive (Motivation); mood or anxiety symptoms (Mood); ability to delay gratification and control behavior, impulses, oral intake and/or changes in reward (Impulsivity); following societal norms and having social graces, tact, and empathy (Conformity); and strongly held beliefs and sensory experiences (Held beliefs). Functional status was evaluated by the Activities of Daily Living Inventory (iADL) questionnaire^22^.

#### Testing for biomarkers

Performed using the Simoa technology on a HD-X analyzer inhouse, using p-tau217 Alzpath and GFAP+NfL Neurology 2-Plex B Assay kits. For p-tau217 we used the formal 0.444pg/ml cutoff, as determined by a recent comparative study^23^.

### APOE genotyping

#### Direct PCR

A sample of patients PBMC DNA (10µl) was heated to 60°C for 10min following incubation on ice. 1µl of sample was used in 15µl PCR reaction with Q5 master mix (NEB, M0592) and the following primers that span the rs429358 SNP site and contain Nextera adapter sequences overhangs: 5’-TCGTCGGCAGCGTCAGATGTGTATAAGAGACAG CACGGCTGTCCAAGGAG-3’ and 5’-GTCTCGTGGGCTCGGAGATGTGTATAAGAGACAGGCGCTTCTGCAGGTCAT-3’ (35 cycles of 98C for 10sec, 67C for 15sec, 72C for 20 sec). Kapa Pure Beads (Kapa KK8002) were used to clean up the amplicons (1x ratio).

#### Library PCR and amplicon sequencing

Library PCR of the amplicons were performed using PrimeStar (Takara, R045) and Nextera UDI primers (8 cycles of: 98°C for 10 sec, 55°C for 10 sec, 72°C for 30 sec). The resulted libraries were pooled and a clean-up with Kapa beads (1x) was performed. The libraries were sequenced on Illumina’s NovaSeq 6000 or NextSeq 2000 platforms. Demultiplexing and fastq files generation was done with Illumina bcl2fastq software and the occurrence of the APOE3/4 alleles was calculated.

### Statistics

#### Biomarker comparison

We compared the correlation between biomarkers using spearman correlations, following previous works methodology^23^. When checking changes in biomarkers distribution between groups (e.g. cognitive healthy vs MCI, smoking vs not smoking), statistical tests were defined separately based on the feature nature in question continues, ordinal or categorical, and fitted an appropriate statistical test Man Whitney U test, for continues and ordinal and chi2 test for categorical features.

When comparing the association between groups over a continuous parameter, referenced as trait associations, we used a linear model corrected for confounders: sex and age.

Correction for multiple hypothesis was done using Benjamini Hochberg FDR correction. All using SciPy ^62^, and code can be found in high_vs_low.py, and trait_association.py.

### Tools for analyses and for creating a predictive model

#### Data preprocess

The preprocess stage included data conversion for numerical representation and normalization as well as feature selection. During the feature selection process, we removed from the data features that were not used in downstream analyses, such as date of p-tau, date of recruitment etc. we excluded samples without basic features such as age, MOCA score, and p-tau217. For categorical features without order relations, we used one-hot-encoding and for categorical features with order relations we used label encoding using the sklearn.preprocess module^63^. Check Preprocess.ipynb for further details. We used a MoCA score threshold of under 26 to label samples as MCI.

We normalized features distribution using sklearn.preprocessing^63^ standard scaler fitting and transforming only on the train set and transforming on all other datasets.

While our cohort is extensive and rich in data, the natural progression of clinical study caused many missing features in the data, for example we started gathering behavioral tests only a few months in the study. We decided to exclude all subjects that did not had all reported features in the INFORMATIVE_FEATURES list in config.py and specified below.

We divided our cohort into two classes: 0 if subject p-tau217 level is below 0.444^23^ and 1 if it’s above. For training and prediction, the model used cognitive features, behavioral features, demographic characteristics, blood tests and selected risk factors. The exact list of features was based on the statistical analysis, choosing the significant features as well as boarder line features

#### Outliers’ removal

We excluded 4 outliers from the cohort while training the prediction model. The subjects are referenced in the main table fight_AD.xlsx as S8, S183, S228, S243 these subjects were excluded for having an extreme value in one or more biomarkers reaching levels of more than twice the value of the second highest samples.

#### Train test split

We used a train-test split of 0.7, 0.3 and when needed split the test set into two equal groups for validation and test sets, resulting in 70:15:15 split. The different datasets were stratified based on the target variable (p-tau217 class) ratio to make sure there isn’t over or under representation of the positive less common class of high p-tau (above 0.444).

#### Bootstrap analysis

We also ran bootstrap analysis using 10 runs with 0.2 train test split to check the robustness of the results, over different metrices – F1, recall, precision, accuracy, AUC, and PRAUC. Check bootstrap_evaluation.py for more details.

#### Comparison metrics

We evaluated the models over an array of metrics such as F1, recall, precision, accuracy, AUC, and PRAUC. We chose the F1 metric to deal with the unbalanced nature of the data for model evaluation.

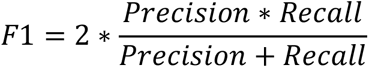

#### Hyperparameters optimization

We used the optuna^64^ package for hyperparameter optimization (HPO); we used different number of trails and optimized catboost^65^, random forest, and logistic regression using different number of trails over different parameters. The metric for optimization was F1 score and was calculated over the validation set. See hyperparameter_optimization.py for further details.

#### Baseline classifier

The baseline classifier was constructed to be similar to the clinical process of the neurologist when choosing which subjects to choose for further testing, for example PET scan or biomarker measurements. To do so we used our MCI threshold of below 27 in the MOCA test score. Check baseline_classifier.py for full implementation.

#### Modeling

We trained different models on this dataset, such as the baseline classifier, catboost^65^, random forest^66^, logistic regression (linear model), and TabPFN2^33^ (tabular foundation model). In the main text we present the results of TabPFN2 model, as it performed best on our evaluation criteria. See the biomarker_prediction directory and tau_classification.py script for full training, evaluation, and benchmarking code.

#### Model evaluation

Due to the unbalanced nature of the data, we evaluated our models based on F1 score, recall, and precision. We also report the model accuracy, AUC and PRAUC to present the full scale of the model performance and limitations.

#### Explainability

In the explainability analysis we used two methods, Accumulated Local Effects (ALE)^34^ using the alibi^67^ package. Check tabpfn_explainability.py for implementation. For explaining the logistic regression model, we used glass box model methodologies. Taking the absolute value over each feature coefficient. Check biomarker_prediction/explaibability directory for code and implementation details.

## Data availability

The data used in the analyses presented in this study are available in the Source Data file. Additional information is available upon reasonable request.

## Code availability

The code used in this study for data analysis, modeling and generation of figures is available at: https://github.com/naomihabiblab/PreclinicalAD

## Acknowledgements

We thank the individuals who volunteered to participate in the study cohort. The work was supported by the Alzheimer’s Foundation of America, and the Israel Science Foundation (ISF) research grant no. 3104/22 (T.B.H. and N.H.). N.H. is supported by the ISF research grant no. 1709/19, the European Research Council (ERC) grant 853409, the Myers Foundation (N.H.), the Chan Zuckerberg Initiative (CZI) Collaborative Pairs grant CP2-1-0000000058. Y.R. is supported by the Beker scholarship for excellence graduate students in cognition and brain science, and the VTAT scholarship for PhD research in data science & AI.

## Contributions

T.B.H. and N.H. designed the study, supervised, and guided the work; L.T. assembled the cohort and all cognitive, medical, and plasma biomarker data, and coordinated the work of A.T., Y.S., F.A., N.N., and Y.L., with input from N.F. and supervision and support from T.B.H.; M.A. guided the experimental design and conducted genetic profiling; Y.R. performed the computational modeling and statistical analyses under the guidance of N.H.; T.B.H., N.H., L.T., and Y.R. wrote the manuscript, and all the co-authors provided critical comments.

## Extended Data Tables & Figures

**Extended Data Table 1:** Clinical characteristics and measurable risk factors of the study cohort. Baseline metabolic, lipid, inflammatory, hematological, blood pressure, and renal parameters for the entire cohort. All physiological and laboratory values are expressed as Mean ± SD alongside their respective normal clinical reference ranges for adults aged 65–80.

| Measurable risk factor | Normal values (ages 65–80) | Mean ± SD (All cohort) |
| --- | --- | --- |
| Metabolic & Diabetes |  |  |
| HbA1c (%) | < 6.1 | 5.70 ± 0.64 |
| Lipid Profile |  |  |
| Total cholesterol (mg/dL) | < 200 | 174.73 ± 40.85 |
| LDL cholesterol (mg/dL) | < 100 | 96.71 ± 34.76 |
| HDL cholesterol (mg/dL) | ≥ 60 | 56.90 ± 15.52 |
| Total cholesterol / HDL ratio | < 3.5 | 3.14 ± 0.88 |
| Triglycerides (mg/dL) | < 150 | 112.28 ± 54.31 |
| Inflammatory & Immune |  |  |
| C-reactive protein (mg/L) | < 1 | 0.93 ± 2.12 |
| White blood cells (10<sup>9</sup>/L) | 4.0–10.0 | 6.62 ± 1.72 |
| Lymphocytes (10<sup>9</sup>/L) | 1.0–4.0 | 2.08 ± 0.74 |
| Hematological Profile |  |  |
| Hemoglobin (g/dL) | 12.0–17.5 | 13.98 ± 1.23 |
| Blood Pressure |  |  |
| Systolic BP (mmHg) | < 130 | 135.21 ± 17.67 |
| Diastolic BP (mmHg) | < 80 | 74.22 ± 10.29 |
| Renal Function |  |  |
| GFR (mL/min/1.73m<sup>2</sup>) | > 72 | 73.49 ± 13.76 |
| Creatinine (mg/dL) | < 1.2 | 0.90 ± 0.13 |

All values are expressed as Mean ± SD unless otherwise specified.

**Extended Data Table 2:**
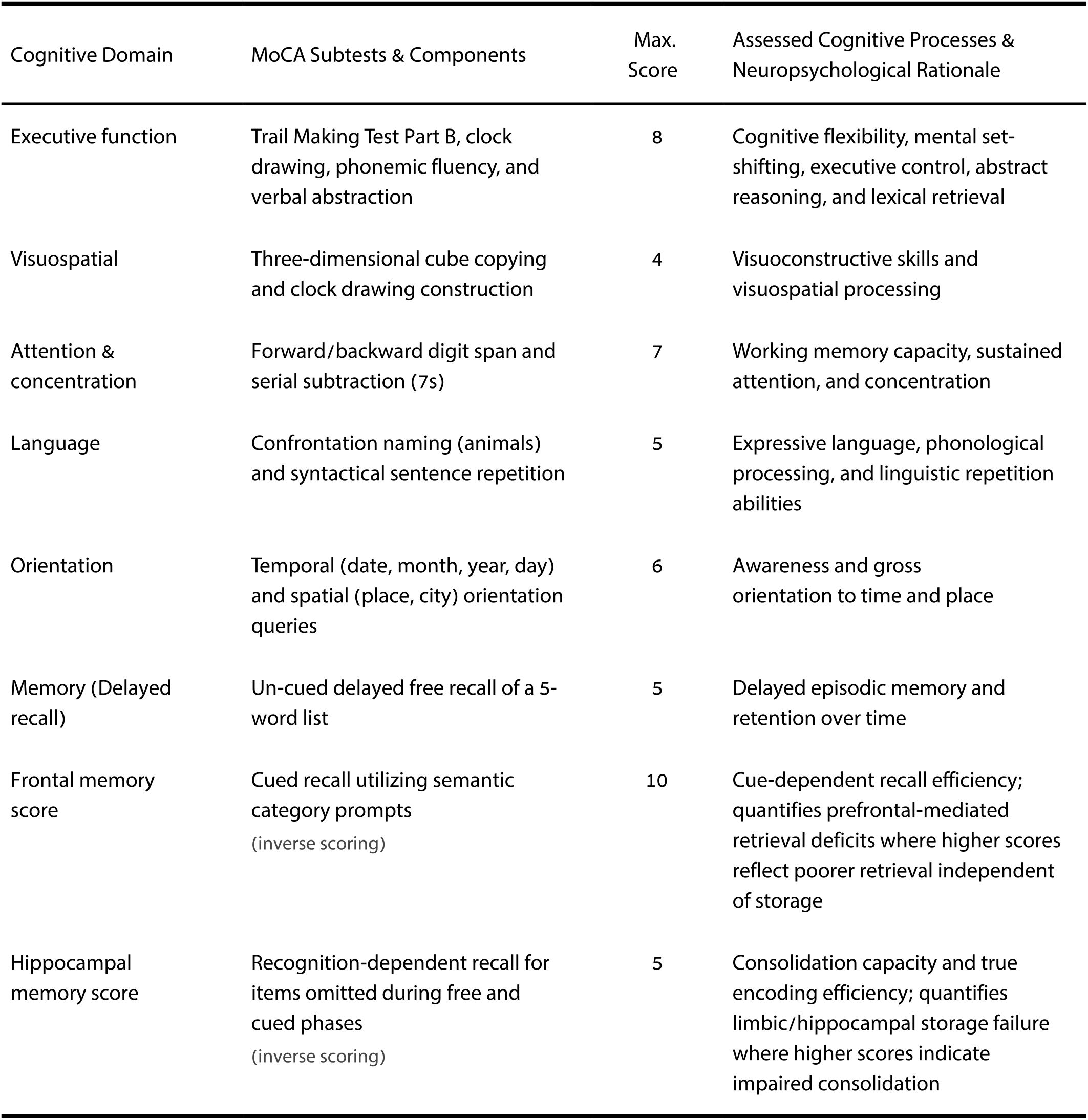
Structural breakdown, scoring parameters, and cognitive processes of MoCA subtests. Details the maximum scores and neuropsychological rationale for each cognitive domain evaluated, including specific tests for executive function, visuospatial processing, attention, language, orientation, and delayed memory. The table additionally outlines the inverse scoring parameters for Frontal memory and Hippocampal memory, which quantify prefrontal-mediated retrieval deficits and limbic/hippocampal consolidation failures, respectively.

**Extended Data Figure 1:**
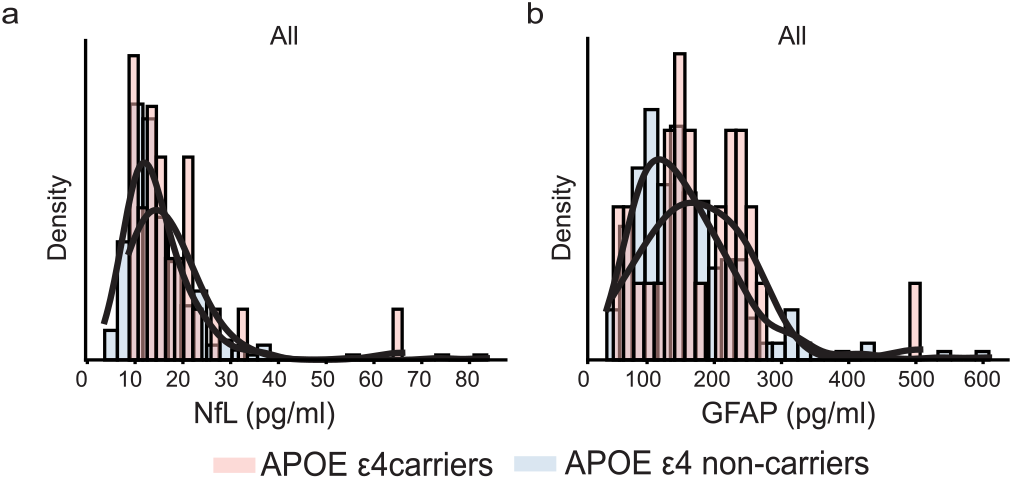
Density distributions of plasma NfL and GFAP stratified by APOE ε4 carrier status. **(a-b)** Density distributions of plasma NfL and GFAP in APOE ε4 carriers and non-carriers across the full cohort (n=251).

**Extended Data Figure 2:**
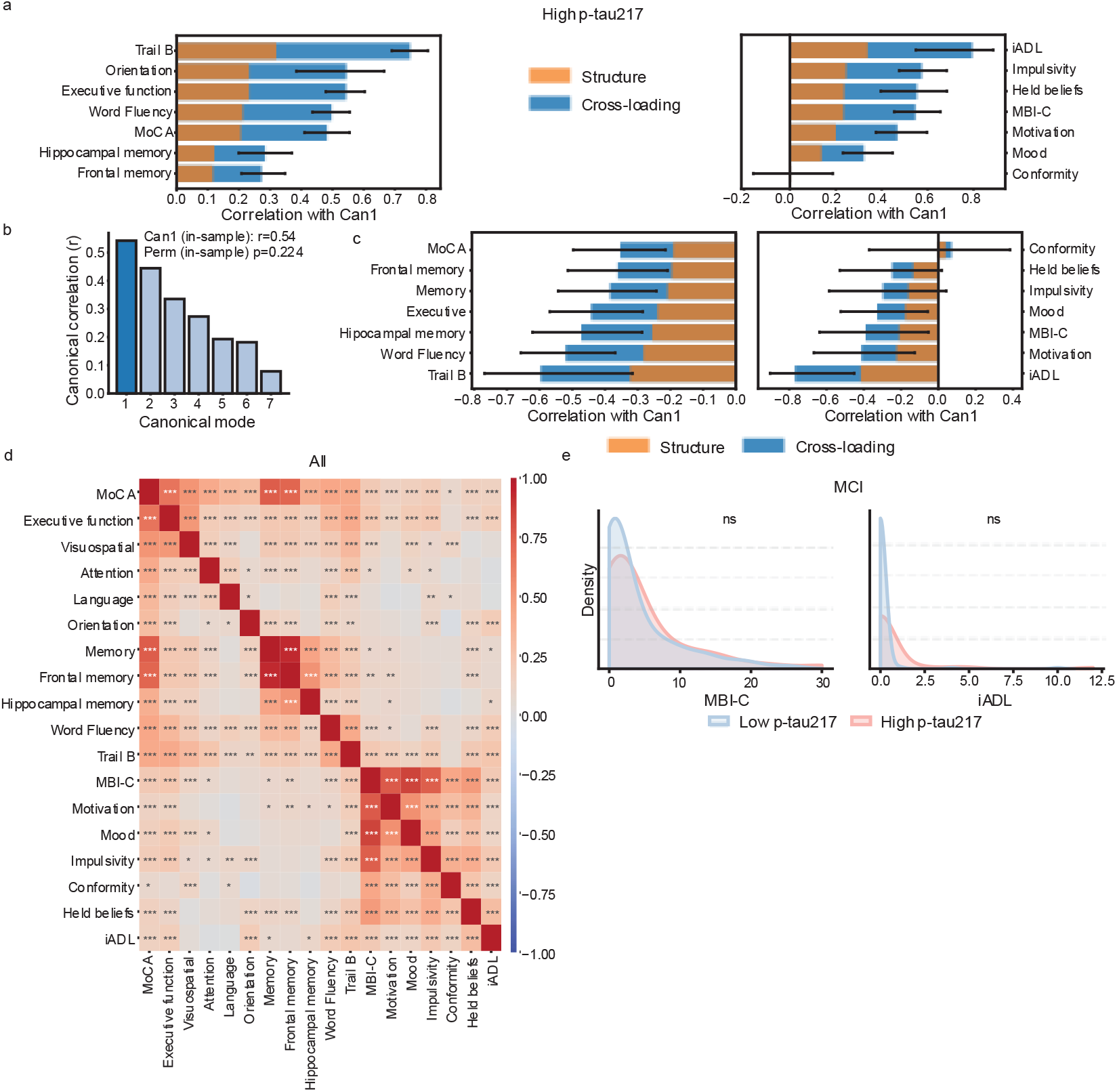
Extended cognitive-behavioral canonical correlations and behavioral density distributions. **(a)** Canonical correlation analysis (CCA) structure and cross-loadings for the full cohort (n=623), demonstrating a similar canonical pattern across the full population. Demonstrating a shared canonical mode linking cognitive (left) and behavioral(right) measures. Cognitive variables represent reductions in performance. Permutation-based p-value test for significance. **(b-c)** structure and cross-loadings for the plasma p-tau217 high subgroup (n=123). First canonical vector (can1) spearman rho=0.54, permutation test (perm) p-value=0.224. Cognitive variables represent reductions in performance. **(d)** Extended Pearson correlation matrix (heatmap) for the entire cohort showing associations between behavioral shifts and the reduction in cognitive scores across domains **(e)** Density distributions of MBI-C and iADL scores stratified by p-tau217 status within the MCI subgroup

**Extended Data Figure 3:**
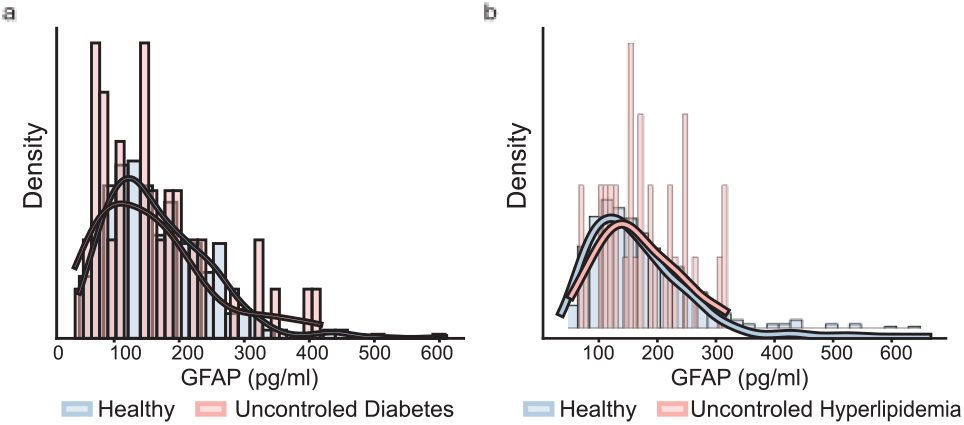
Metabolic risk-factor control stratified plasma GFAP levels biomarker. **(a-b)** Density distributions of plasma GFAP according to control of metabolic risk-factors diabetes **(a)** and hyperlipidemia **(b)**, showing no association between risk control and GFAP levels.

**Extended Data Figure 4:**
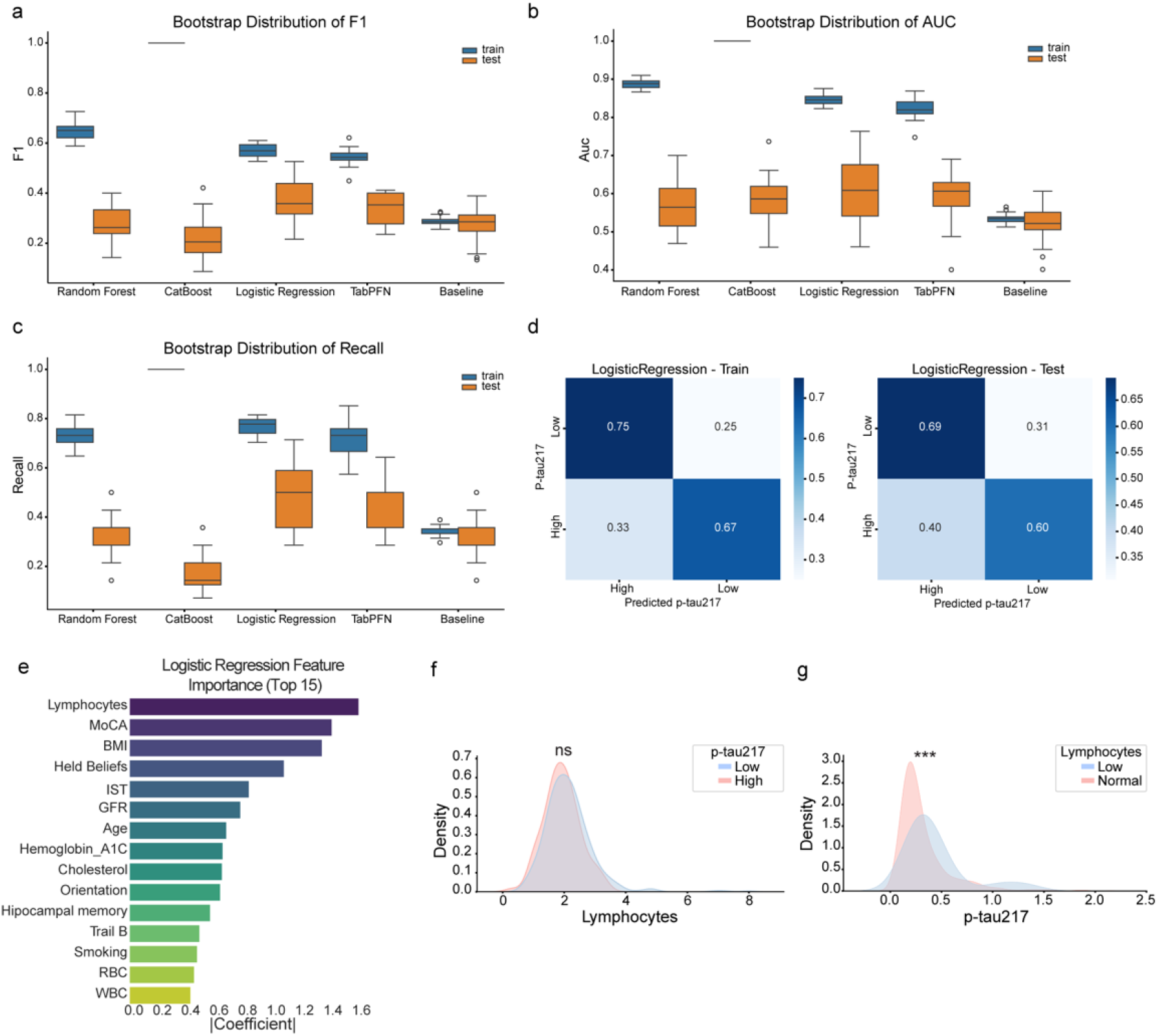
Comparative evaluation and interpretation of machine-learning models for p-tau217 prediction. **(a-c)** Bootstrap distributions of model performance metrics across training and test datasets (train-test split 85:15). Comparing the baseline classifier, Random Forest, CatBoost, Logistic Regression, and TabPFN models. Showing F1 score **(a)**, Area under the receiver operating characteristic curve (AUC) **(b)**, and recall **(c)**. **(d)** Training and test confusion matrices for the logistic regression model classifier, normalized by row. **(e)** Top 15 logistic regression features ranked by the absolute coefficient magnitude. **(f)** Distribution of lymphocyte counts stratified by plasma p-tau217 status, demonstrating no significant difference in the distribution. **(g)** Distribution of plasma p-tau217 levels comparing low lymphocyte count (<1×10⁹ /L) versus normal counts, illustrating increased prevalence of lymphopenia at higher p-tau217 levels, in alignment with the predictive value of the lymphocyte count feature in the classifier.

## References

1. Vermunt, L. et al. Duration of preclinical, prodromal, and dementia stages of Alzheimer’s disease in relation to age, sex, and APOE genotype. Alzheimers Dement 15, 888–898 (2019).

2. Jia, J. et al. Biomarker Changes during 20 Years Preceding Alzheimer’s Disease. N Engl J Med 390, 712–722 (2024).

3. Zheng, Q. & Wang, X. Alzheimer’s disease: insights into pathology, molecular mechanisms, and therapy. Protein Cell 16, 83–120 (2025).

4. Kusoro, O., Roche, M., Del-Pino-Casado, R., Leung, P. & Orgeta, V. Time to Diagnosis in Dementia: A Systematic Review With Meta-Analysis. Int J Geriatr Psychiatry 40, e70129 (2025).

5. Ossenkoppele, R. et al. Plasma p-tau217 and tau-PET predict future cognitive decline among cognitively unimpaired individuals: implications for clinical trials. Nat Aging 5, 883–896 (2025).

6. Plasma phosphorylated tau 217 and longitudinal trajectories of Aβ, tau, and cognition in cognitively unimpaired older adults. Nature Communications. https://www.nature.com/articles/s41467-026-71269-3.

7. Buckley, R. F. et al. Prognostic Value of Blood-Based P-Tau217 Levels for Progression to Cognitive Impairment. JAMA e2612556 (2026) doi:10.1001/jama.2026.12556.

8. Wang, H. et al. Biomarkers associated with the pathogenesis of Alzheimer’s disease. Front Cell Neurosci 17, 1279046 (2023).

9. Kapasi, A., DeCarli, C. & Schneider, J. A. Impact of multiple pathologies on the threshold for clinically overt dementia. Acta Neuropathol 134, 171–186 (2017).

10. Leffa, D. T. et al. In vivo-measured Lewy body pathology is associated with neuropsychiatric symptoms across the Alzheimer’s disease continuum. Mol Psychiatry 31, 2472–2480 (2026).

11. Caselli, R. J. et al. Neuropsychological decline up to 20 years before incident mild cognitive impairment. Alzheimers Dement 16, 512–523 (2020).

12. Livingston, G., et al. Dementia prevention, intervention, and care: 2024 report of the Lancet standing Commission. Lancet 404, 572–628 (2024).

13. Nichols, E. et al. Differences in the prevalence and patterns of dementia risk factors across 14 countries and regions: a harmonised cross-national analysis. The Lancet Healthy Longevity 0, (2026).

14. Nedelec, T. et al. Identifying health conditions associated with Alzheimer’s disease up to 15 years before diagnosis: an agnostic study of French and British health records. The Lancet Digital Health 4, e169–e178 (2022).

15. Stephan, B. C. M. et al. Population attributable fractions of modifiable risk factors for dementia: a systematic review and meta-analysis. The Lancet Healthy Longevity 5, e406–e421 (2024).

16. Roccati, E. et al. Modifiable dementia risk factors and AT(N) biomarkers: findings from the EPAD cohort. Front. Aging Neurosci. 16, (2024).

17. Wang, J., Gu, B. J., Masters, C. L. & Wang, Y.-J. A systemic view of Alzheimer disease - insights from amyloid-β metabolism beyond the brain. Nat Rev Neurol 13, 612–623 (2017).

18. Nasreddine, Z. S. et al. The Montreal Cognitive Assessment, MoCA: a brief screening tool for mild cognitive impairment. J Am Geriatr Soc 53, 695–699 (2005).

19. Tombaugh, T. N. Trail Making Test A and B: normative data stratified by age and education. Arch Clin Neuropsychol 19, 203–214 (2004).

20. Isaacs, B. & Akhtar, A. J. The set test: a rapid test of mental function in old people. Age Ageing 1, 222–226 (1972).

21. Ismail, Z. et al. The Mild Behavioral Impairment Checklist (MBI-C): A Rating Scale for Neuropsychiatric Symptoms in Pre-Dementia Populations. J Alzheimers Dis 56, 929–938 (2017).

22. Lawton, M. P. & Brody, E. M. Assessment of older people: self-maintaining and instrumental activities of daily living. Gerontologist 9, 179–186 (1969).

23. Schindler, S. E. et al. Head-to-head comparison of leading blood tests for Alzheimer’s disease pathology. Alzheimers Dement 20, 8074–8096 (2024).

24. Charach, G. et al. Apolipoprotein E (APOE) Genotypes in the Israeli population. Isr Med Assoc J 25, 803–807 (2022).

25. Villemagne, V. L. et al. Amyloid β deposition, neurodegeneration, and cognitive decline in sporadic Alzheimer’s disease: a prospective cohort study. Lancet Neurol 12, 357–367 (2013).

26. Milà-Alomà, M. et al. Amyloid beta, tau, synaptic, neurodegeneration, and glial biomarkers in the preclinical stage of the Alzheimer’s continuum. Alzheimers Dement 16, 1358–1371 (2020).

27. Benedet, A. L. et al. Differences Between Plasma and Cerebrospinal Fluid Glial Fibrillary Acidic Protein Levels Across the Alzheimer Disease Continuum. JAMA Neurol 78, 1471–1483 (2021).

28. Simons, J. S., Garrison, J. R. & Johnson, M. K. Brain Mechanisms of Reality Monitoring. Trends Cogn Sci 21, 462–473 (2017).

29. Moran, C. et al. Brain atrophy in type 2 diabetes: regional distribution and influence on cognition. Diabetes Care 36, 4036–4042 (2013).

30. Abner, E. L. et al. Diabetes is associated with cerebrovascular but not Alzheimer’s disease neuropathology. Alzheimer’s & Dementia 12, 882–889 (2016).

31. Wingo, A. P. et al. LDL cholesterol is associated with higher AD neuropathology burden independent of APOE. J Neurol Neurosurg Psychiatry 93, 930–938 (2022).

32. Eyting, M. et al. A natural experiment on the effect of herpes zoster vaccination on dementia. Nature 641, 438–446 (2025).

33. Grinsztajn, L., et al. TabPFN-2.5: Advancing the State of the Art in Tabular Foundation Models. arXiv.org https://arxiv.org/abs/2511.08667v2 (2025).

34. Visualizing the Effects of Predictor Variables in Black Box Supervised Learning Models. Journal of the Royal Statistical Society Series B: Statistical Methodology | Oxford Academic. https://academic.oup.com/jrsssb/article/82/4/1059/7056085?login=true.

35. Twamley, E. W., Ropacki, S. A. L. & Bondi, M. W. Neuropsychological and neuroimaging changes in preclinical Alzheimer’s disease. J Int Neuropsychol Soc 12, 707–735 (2006).

36. Grober, E. et al. Memory impairment, executive dysfunction, and intellectual decline in preclinical Alzheimer’s disease. J Int Neuropsychol Soc 14, 266–278 (2008).

37. Wilson, R. S. et al. Neuropsychiatric symptoms in Brazilians with mild cognitive impairment and dementia. Alzheimers Dement (Amst) 13, e12219 (2021).

38. Lyketsos, C. G. & Peters, M. E. Neuropsychiatric symptoms in Alzheimer’s disease: Past, present, and future. Alzheimers Dement 22, e71077 (2026).

39. Chen, Y., Dang, M. & Zhang, Z. Brain mechanisms underlying neuropsychiatric symptoms in Alzheimer’s disease: a systematic review of symptom-general and –specific lesion patterns. Mol Neurodegener 16, 38 (2021).

40. Alves, L. et al. Neuropsychological profile of amyloid-positive versus amyloid-negative amnestic Mild Cognitive Impairment. J Neuropsychol 15 Suppl 1, 41–52 (2021).

41. Asken, B. M. et al. Informing etiological heterogeneity of mild cognitive impairment and risk for progression to dementia with plasma p-tau217. The Journal of Prevention of Alzheimer’s Disease 12, 100011 (2025).

42. Malpetti, M. et al. Regional tau PET patterns predict prospective domain-specific cognitive decline in early symptomatic Alzheimer’s disease. Alzheimers Res Ther 17, 220 (2025).

43. Livingston, G., et al. Dementia prevention, intervention, and care: 2024 report of the Lancet standing Commission. The Lancet 404, 572–628 (2024).

44. The SPRINT MIND Investigators for the SPRINT Research Group. Effect of Intensive vs Standard Blood Pressure Control on Probable Dementia: A Randomized Clinical Trial. JAMA 321, 553–561 (2019).

45. Ngandu, T. et al. A 2 year multidomain intervention of diet, exercise, cognitive training, and vascular risk monitoring versus control to prevent cognitive decline in at-risk elderly people (FINGER): a randomised controlled trial. Lancet 385, 2255–2263 (2015).

46. Pappolla, M. A., Refolo, L., Sambamurti, K., Zambon, D. & Duff, K. Hypercholesterolemia and Alzheimer’s Disease: Unraveling the Connection and Assessing the Efficacy of Lipid-Lowering Therapies. J Alzheimers Dis 101, S371–S393 (2024).

47. Ahmed, H. et al. Brain cholesterol and Alzheimer’s disease: challenges and opportunities in probe and drug development. Brain 147, 1622–1635 (2024).

48. Frison, E. et al. Diabetes Mellitus and Cognition: Pathway Analysis in the MEMENTO Cohort. Neurology 97, e836–e848 (2021).

49. Moran, C. et al. The Association of Type 2 Diabetes Mellitus with Cerebral Gray Matter Volume Is Independent of Retinal Vascular Architecture and Retinopathy. J Diabetes Res 2016, 6328953 (2016).

50. Croese, T. et al. Disruption of the brain-spleen axis impairs monocyte-microglia communication and accelerates disease progression in a mouse model of amyloidosis. Nat Commun https://doi.org/10.1038/s41467-026-74253-z (2026) doi:10.1038/s41467-026-74253-z.

51. Wang, S. et al. Association of impaired kidney function with dementia and brain pathologies: A community-based cohort study. Alzheimers Dement 19, 2765–2773 (2023).

52. Guo, J. et al. Body Mass Index Trajectories Preceding Incident Mild Cognitive Impairment and Dementia. JAMA Psychiatry 79, 1180–1187 (2022).

53. Dang, M. et al. Stage-specific temporal associations between body mass index trajectories and Alzheimer’s disease pathologies. Eur Radiol 36, 4891–4903 (2026).

54. Arnold, D. et al. Stratifying dementia risk factors: A prediction model and hypothesis-driven analysis. Alzheimers Dement 21, e70870 (2025).

55. Arslan, B. et al. The impact of kidney function on Alzheimer’s disease blood biomarkers: implications for predicting amyloid-β positivity. Alz Res Therapy 17, 48 (2025).

56. Mondésert, E. et al. Influence of Decreased Kidney Function on Plasma Biomarkers of Neurodegenerative Disorders in Routine Care: Confirmation of the Interest of Ratios. Neurology 106, e214931 (2026).

57. Morris, J. K., Honea, R. A., Vidoni, E. D., Swerdlow, R. H. & Burns, J. M. Is Alzheimer’s disease a systemic disease? Biochim Biophys Acta 1842, 1340–1349 (2014).

58. Heneka, M. T. et al. Neuroinflammation in Alzheimer’s disease. Lancet Neurol 14, 388–405 (2015).

59. Cai, H. et al. Systemic inflammatory markers in ageing, Alzheimer’s disease and other dementias. Brain 148, 480–492 (2025).

60. Ye, Z. et al. Leveraging routine clinical data for dementia risk prediction using machine learning. Journal of Alzheimer’s Disease 111, 1516–1526 (2026).

61. Park, J. H. et al. Machine learning prediction of incidence of Alzheimer’s disease using large-scale administrative health data. npj Digit. Med. 3, 46 (2020).

62. Virtanen, P. et al. SciPy 1.0: fundamental algorithms for scientific computing in Python. Nat Methods 17, 261–272 (2020).

63. Pedregosa, F. et al. Scikit-learn: Machine Learning in Python. Journal of Machine Learning Research 12, 2825–2830 (2011).

64. Optuna - A hyperparameter optimization framework. Optuna https://optuna.org/.

65. Prokhorenkova, L., Gusev, G., Vorobev, A., Dorogush, A. V. & Gulin, A. CatBoost: unbiased boosting with categorical features. Preprint at 10.48550/arXiv.1706.09516 (2019).

66. Breiman, L. Random Forests. Machine Learning 45, 5–32 (2001).

67. Klaise, J., Looveren, A. V., Vacanti, G. & Coca, A. Alibi Explain: Algorithms for Explaining Machine Learning Models. Journal of Machine Learning Research 22, 1–7 (2021).

